# VERSE: a versatile and efficient RNA-Seq read counting tool

**DOI:** 10.1101/053306

**Authors:** Qin Zhu, Stephen A Fisher, Jamie Shallcross, Junhyong Kim

## Abstract

**Motivation:** RNA-Seq is a powerful technology that delivers digital gene expression data. To measure expression strength at the gene level, one popular approach is direct read counting after aligning the reads to a reference genome/transcriptome. HTSeq is one of the most popular ways of counting reads, yet its slow running speed of poses a bottleneck to many RNA-Seq pipelines. Gene level counting programs also lack a robust scheme for quantifying reads that map to non-exonic genomic features, such as intronic and intergenic regions, even though these reads are prevalent in most RNA-Seq data.

**Results:** In this paper we present VERSE, an RNA-Seq read counting tool which builds upon the speed of featureCounts and implements the counting modes of HTSeq. VERSE is more than 30x faster than HTSeq when computing the same gene counts. VERSE also supports a hierarchical assignment scheme, which allows reads to be assigned uniquely and sequentially to different types of features according to user-defined priorities.

**Availability:** VERSE is implemented in C. It is built on top of featureCounts. VERSE is open source and can be downloaded freely from Github (https://github.com/qinzhu/VERSE).

**Supplementary information:** Tables and figures illustrating the counting modes implemented in VERSE and the differences between hierarchical and independent assignment.

## 1 Introduction

Turning raw RNA sequencing data into a meaningful gene expression table requires expression quantification, which has substantial influence on the quality of downstream analyses (Fonseca et al., 2014; Oshlack et al., 2010). Current popular expression quantification tools can be roughly grouped into two categories: tools such as HTSeq (Anders et al., 2014), featureCounts (Liao et al., 2013), Rcount (Schmid and Grossniklaus, 2014) and summarizeOverlaps (Lawrence et al., 2013) that compute expression by *directly counting* the number of reads overlapping features, and tools such as Cufflinks (Trapnell et al., 2012), RSEM (Li and Dewey, 2011) and EMSAR (Lee et al., 2015) that compute expression by *estimating* abundance at the transcript level. Here we focus on tools that directly count expression based on genomic mappings. These tools define genomic coordinate intervals of features and count the number of reads that fall within each interval. Complications arise because each feature may comprise multiple genomic coordinate intervals, two or more features may overlap in their intervals, or the reads may map partially outside of feature intervals as well as to multiple genomic coordinates (Supplementary Table 1). The popular program HTSeq implements three different counting modes to address these complications, but the original python implementation does not provide satisfactory running speed for high-throughput data analysis.

An important aspect of read counting is to quantify not only exons, but the full spectrum of feature types that may produce RNA transcripts. It is often found that intronic and intergenic reads make up a significant proportion of RNA-Seq data (Oshlack et al., 2010). These reads may represent genomic contamination, or authentic biological transcripts such as pre-mRNA or non-coding RNA (Kapranov et al., 2011). Current available read assignment tools only allow one feature type to be quantified at one time, forcing users to go through significant effort and even slower pipeline processing to capture read assignments to multiple feature types. There are also many complications in handling reads located at the boundary or overlapping region of different feature types. The proper handling of read assignments across feature types is becoming even more complex as RNA-Seq techniques are being applied to human samples. For example, the GENCODE human reference gene set has three levels of genes, representing experimental validated, manually annotated and algorithm predicted transcripts respectively (Harrow et al., 2012), each with different degree of overlap. To our knowledge, none of the current available read counting tools is capable of directly assigning reads to all three annotation levels without over-counting.

We developed VERSE with the goal of very fast read assignments with flexible counting modes. Additionally, VERSE provides a hierarchical assignment scheme which allows reads to be quantified sequentially to different feature types or annotation levels in a single run.

## 2 Description

VERSE supports all three HTSeq read assignment modes, a modified intersection-nonempty mode, a mode that assigns reads to features based on overlapping length and the featureCount mode. Unlike the genomic array method utilized by HTSeq, VERSE uses an end-comparison method to implement the set operation process. This method, combined with the feature block data structure of featureCounts, allows VERSE to process reads at a much faster speed while outputting the exact HTSeq-style count result. We compared the performance of HTSeq, summarizeOverlaps and VERSE using data from Dueck et al (Dueck et al., 2015) and the mm10 (Dec. 2011 assembly) annotation file obtained from UCSC genome browser (Karolchik, 2003). As shown in Table 1, VERSE is more than 30 times faster than HTSeq and summarizeOverlaps, and multithreading further increased its performance with a slight memory tradeoff.

**Table 1.**
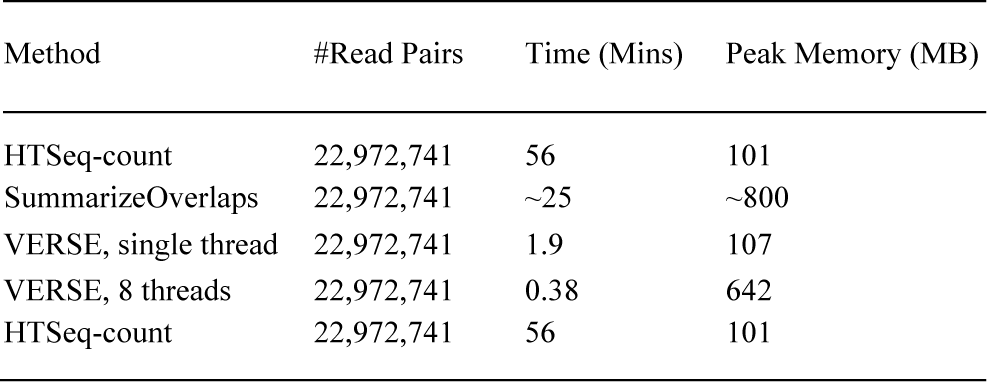
Performance results on the RNA-Seq data of 10 pg total RNA from mouse heart tissue

| Method | #Read Pairs | Time (Mins) | Peak Memory (MB) |
| --- | --- | --- | --- |
| HTSeq-count | 22,972,741 | 56 | 101 |
| SummarizeOverlaps | 22,972,741 | ~25 | ~800 |
| VERSE, single thread | 22,972,741 | 1.9 | 107 |
| VERSE, 8 threads | 22,972,741 | 0.38 | 642 |
| HTSeq-count | 22,972,741 | 56 | 101 |

Note: Only annotated exons were quantified using intersection-nonempty mode. VERSE is faster than both HTSeq and SummarizeOverlaps by two orders of magnitude. The testing results were generated using VERSE 0.1.5, HTSeq-0.6.1p2 and GenomicRanges 1.22.3, on a computer node with 8 Intel(R) Xeon(R) 3.3GHz CPU and 16 GB of memory.

The hierarchical assignment scheme of VERSE supports read assignment over multiple feature types or annotation levels. Figure 1 illustrates the general structure of this scheme. In the hierarchical mode, reads are sequentially assigned to every feature types with the removal of assigned and ambiguous reads at each level. With the above dataset, it only took VERSE 0.45 minutes to complete seven feature type (exon, antisense-exon, mitochondria, antisense-mitochondria, intron, antisense-intron and intergenic) assignments with 8 threads, compared to seven separate runs and more than 5 hours using HTSeq. With HTSeq, approximately 6% of the reads were over-counted, since there’s no way to remove assigned and ambiguous reads from the BAM file, leading to an inaccurate abundance estimation of non-exonic features (Supplementary Figure 1).

**Fig. 1.**
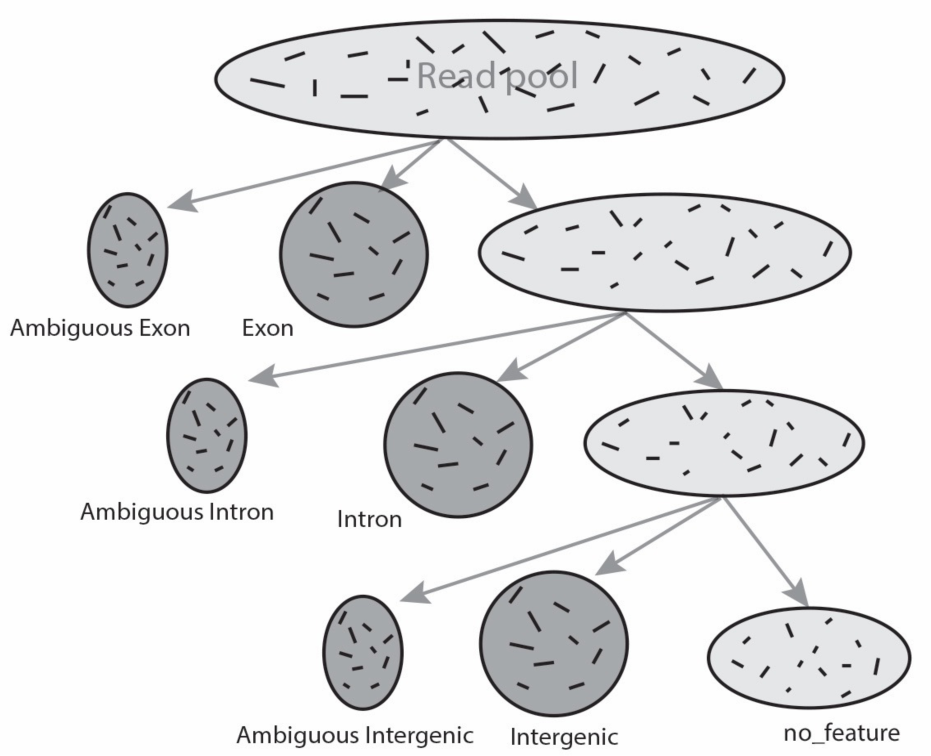
Illustration of hierarchical assignment scheme. Reads are first assigned to the feature type with the highest priority, in this case, exon. Any reads that cannot be matched to exon will be compared with the next feature type annotation, intron. This procedure is repeated until all reads are successfully assigned, set as ambiguous or cannot be mapped to any of the annotations.

With the hierarchical scheme and the modified intersection-nonempty mode, only reads that truly do not map to any feature will descend along the hierarchy. The read assignment hierarchy should be carefully designed so that it does not violate basic biological assumptions while addressing the main research interest. For example, exons are expected to have higher frequency than antisense/intron/intergenic in an mRNA-Seq experiment and thus should be listed first in the hierarchy. Similarly users will typically want level 1 and 2 transcripts in the GENCODE annotation to take priority over level 3 transcripts. The feature types at lower levels of the assignment hierarchy will likely receive fewer read counts, but the false positive rate is also expected to drop significantly, which may be a desirable behaviour in the context of novel transcript discovery.

## Acknowledgements

VERSE is developed based on the framework of featureCounts written by Drs. Yang Liao and Wei Shi.

